# Quercetin alleviates intestinal oxidative stress in Jilin White geese

**DOI:** 10.1101/2023.08.14.553261

**Authors:** Liu Yingkun, Guo Wei, Jia Fangyuan, Wang Kai, Li Zhichao, Zhang Tao, Liu Mo, Zhou Haizhu

## Abstract

Oxidative stress can occur during all stages of geese breeding, frequently damaging the intestinal tract and various tissues and organs. Quercetin is a flavonoid that exerts potential therapeutic effects against oxidative stress-induced cell damage and mitochondrial dysfunction. Herein, we investigated the alleviating effects of quercetin against lipopolysaccharide (**LPS**)-induced intestinal inflammation and oxidative stress in Jilin White geese. We selected 120 healthy male 7-d-old geese of similar BW and randomly divided them into six treatment groups, with five replicates each and four geese in each replicate. The groups were as follows: C (fed basal diet, i.p. injection of 0.5 mg/kg normal saline); L (fed basal diet, i.p. injection of 0.5 mg/kg LPS); Q1, Q2, Q3, and Q4 (diet + 150, 300, 450, or 600 mg/kg quercetin, respectively, i.p. injection of 0.5 mg/kg LPS). Samples were collected after experimental d 21. Our results showed that adding 300 mg/kg quercetin to the diet significantly improved growth performance after oxidative stress and reduced serum low-density lipoprotein and glutathione aminotransferase activities. Adding 450 mg/kg quercetin significantly increased the intestinal expression of downstream genes of the nuclear factor E2-related factor 2 pathway, heme oxygenase 1 (***HO-1***), reduced coenzyme/quinone oxidoreductase (***NQO1***), and glutathione oxidase (***GSTP***) in Jilin White geese. Considering production, adding 300 mg/kg quercetin to the diet could alleviate LPS-induced oxidative stress and reduce costs. These findings provide a reference hypothesis for the oxidative stress damage during geese production, which may help reduce the economic loss owing to geese mortality.

## Introduction

China has the highest population of geese worldwide and the most abundant geese breeding resources. However, artificial selection pressures in intense animal breeding systems and environmental factors can induce oxidative stress, resulting in disease or death and seriously affecting the development of the geese breeding industry [1]. Oxidative stress increases cell membrane permeability, resulting in a series of biochemical abnormalities, such as cytoplasmic membrane rupture, mitochondrial swelling, and expansion of the endoplasmic reticulum [2]. The intestine is an important secretion and immune organ that is sensitive to oxidative stress, which can cause intestinal mucosal damage and increased permeability. This can not only affect the digestion and absorption of nutrients but also lead to intestinal diseases and systemic immunosuppression. In geese, oxidative stress can seriously impact performance and meat quality, resulting in economic losses in the breeding industry[3].Quercetin, a naturally occurring flavonoid with a wide range of biological activities and pharmacological effects, can scavenge free radicals. Quercetin has shown potential therapeutic effects against oxidative stress-induced cell damage and mitochondrial dysfunction [4,5]. Quercetin alleviates iron overload-induced osteoporosis and prevents radiation-induced intestinal oxidative stress through the nuclear factor E2-related factor 2 (**Nrf2**) signaling pathway [6,7]. Nrf2 is a transcription factor involved in defense against a variety of harmful stresses and can improve oxidative stress, as well as help maintain cellular redox homeostasis [8,9]. Nrf2 can regulate downstream antioxidant genes to exert potent antioxidant effects [10]. When the redox state is imbalanced, Nrf2 dissociates from the isolation complex and translocates to the cell to interact with the antioxidant response element of antioxidant genes, causing transcriptional activation of downstream target genes, such as heme oxygenase 1 (***HO-1***), reduced coenzyme/quinone oxidoreductase (***NQO1***), and glutathione oxidase (***GSTP***), to scavenge excess reactive oxygen species [11,12].

In the present study, Jilin White geese were used as test animals to establish a lipopolysaccharide (**LPS**) oxidative stress model. By undertaking feeding experiments, we examined the effects of dietary quercetin addition on the intestinal tract of geese under oxidative stress to elucidate the protective effect of quercetin.

## Materials and methods

### Experimental design

The experimental protocol was approved by the Experimental Ethics Committee of Jilin Agricultural University. All operations were performed under pentobarbital sodium anesthesia, with every effort made to reduce pain. In total, 120 healthy male 7-d-old Jilin White geese (Yuhong Ecological Agriculture Technology Co., Jilin Province, CHN) with similar BW (900 ± 50 g) were randomly allotted into six groups with five replicates per group (n=4 geese/replicate). The groups were as follows: C (basal diet, i.p. injection of 0.5 mg/kg normal saline); stress group was indicated as L (basal diet, i.p. injection of 0.5 mg/kg LPS); treatment groups indicated as Q1, Q2, Q3, and Q4 (diet + 150, 300, 450, or 600 mg/kg quercetin, respectively, i.p. injection of 0.5 mg/kg LPS). Geese were administered LPS injections on experimental d 16 and 21 to induce oxidative stress. LPS was purchased from Sigma Chemical Biotechnology (Louis, Missouri, USA). Quercetin (purity: 99%) was purchased from Shanghai Maclean Biotechnology (Shanghai, CHN). The diets were formulated according to the NRC (2007) feeding standards. Table 1 summarizes the composition and nutritional levels of the experimental basal diet. The geese were raised in 30 separate floor pens, with free access to food and water.

**Table 1.**
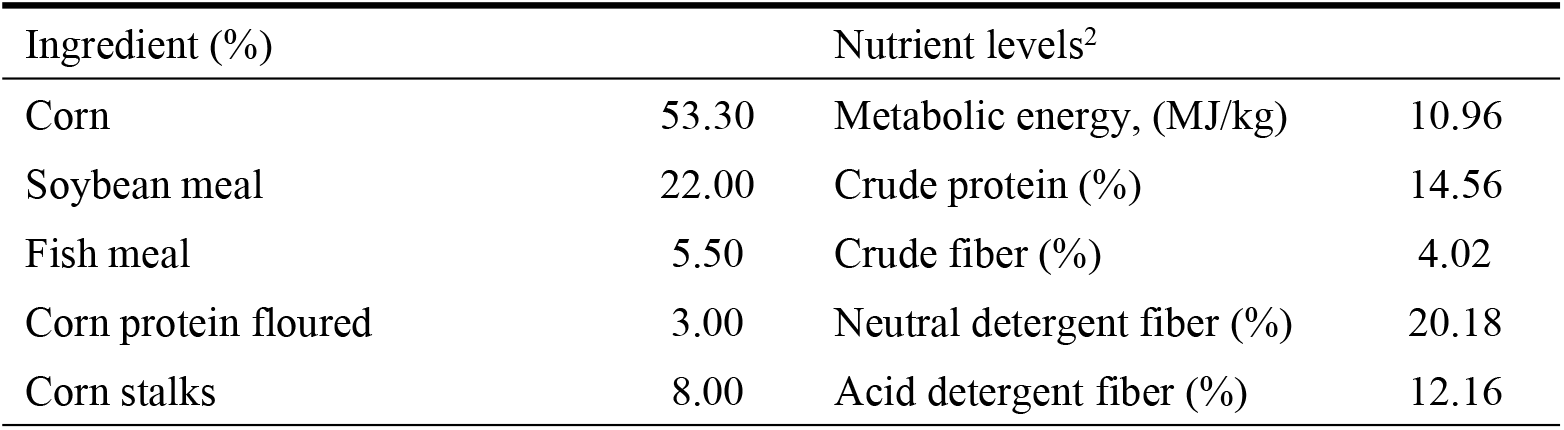
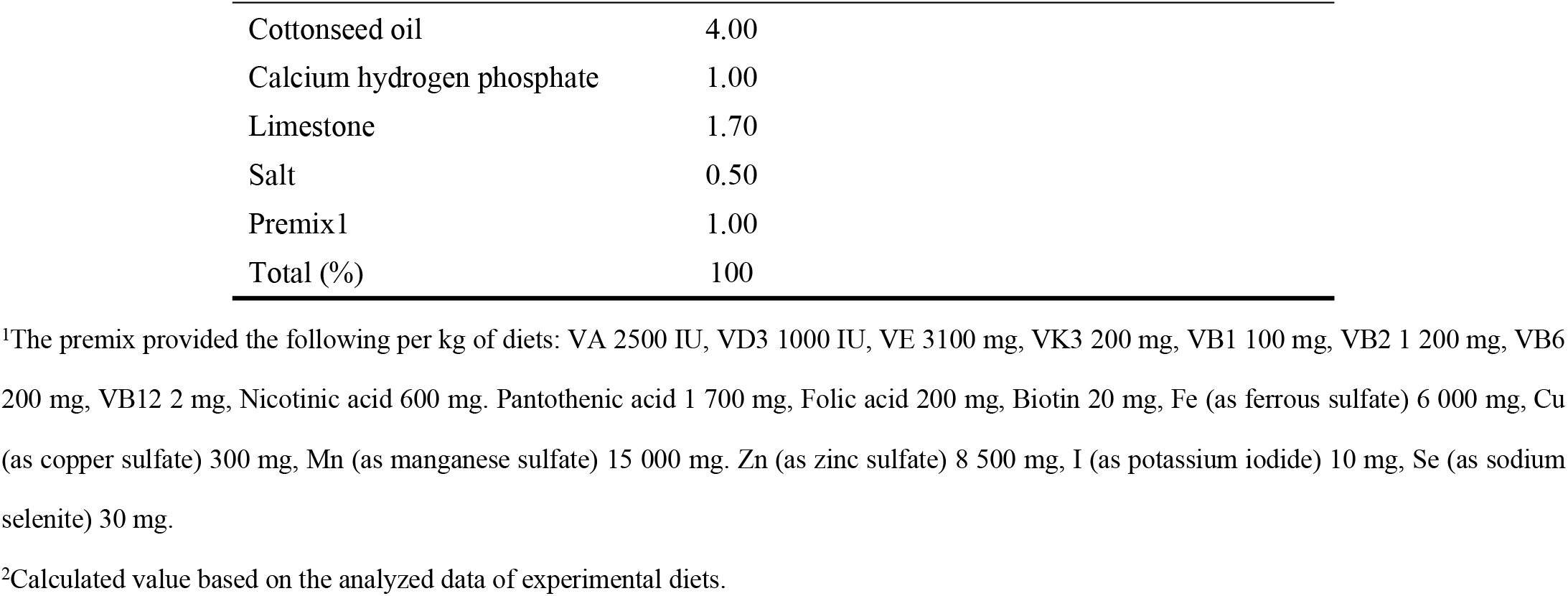
Composition and nutrient levels of the experimental basal diet.

| Ingredient (%) |  | Nutrient levels<sup>2</sup> |  |
| --- | --- | --- | --- |
| Corn | 53.30 | Metabolic energy, (MJ/kg) | 10.96 |
| Soybean meal | 22.00 | Crude protein (%) | 14.56 |
| Fish meal | 5.50 | Crude fiber (%) | 4.02 |
| Corn protein floured | 3.00 | Neutral detergent fiber (%) | 20.18 |
| Corn stalks | 8.00 | Acid detergent fiber (%) | 12.16 |

|  |  |
| --- | --- |
| Cottonseed oil | 4.00 |
| Calcium hydrogen phosphate | 1.00 |
| Limestone | 1.70 |
| Salt | 0.50 |
| Premix 1 | 1.00 |
| Total (%) | 100 |

<sup>1</sup>The premix provided the following per kg of diets: VA 2500 IU, VD3 1000 IU, VE 3100 mg, VK3 200 mg, VB1 100 mg, VB2 1 200 mg, VB6 200 mg, VB12 2 mg, Nicotinic acid 600 mg, Pantothenic acid 1 700 mg, Folic acid 200 mg, Biotin 20 mg, Fe (as ferrous sulfate) 6 000 mg, Cu (as copper sulfate) 300 mg, Mn (as manganese sulfate) 15 000 mg, Zn (as zinc sulfate) 8 500 mg, I (as potassium iodide) 10 mg, Se (as sodium selenite) 30 mg.
<sup>2</sup>Calculated value based on the analyzed data of experimental diets.

### Sample collection and indicator determination

#### Sample Collection

At the end of treatment, 30 geese (one goose per replicate) were randomly selected and feed-deprived for 12 h. The heart, liver, spleen, kidney, and pectoral and leg muscles were harvested and weighed. Blood samples were collected from the wing veins, and geese were then slaughtered by jugular vein exsanguination. The blood samples were centrifuged at 3000 r/min for 10 min at 4°C, and the serum was collected and stored at -80°C. The whole intestinal tract was removed quickly. The tissues of the middle part of the duodenum, jejunum, and ileum were cut using sterile forceps and scissors, rinsed gently with precooled PBS, quickly frozen in liquid nitrogen, and stored at 80°C for mRNA expression analysis.

### Growth performance

During the experiment, feed consumption was recorded daily to determine the average daily feed intake (**ADFI**), initial BW (**IBW**), final BW (**FBW**), average daily gain (**ADG**), and feed conversion ratio (**FCR**). The number of deaths was recorded daily and used to adjust the total number of geese per replicate to exclude them from the calculation of ADFI and FCR.

### Blood Physiological and Biochemical Parameters and Antioxidant Capacity Indicators

The following parameters were measured: concentrations of total cholesterol (**T-CHO**), low-density lipoprotein cholesterol (**LDL-C**), glutathione aminotransferase (**AST**), alanine aminotransferase (**ALT**), alkaline phosphatase (**AKP**), and high-density lipoprotein cholesterol (**HDL-C**); levels of malondialdehyde (**MDA**), 8-hydroxydeoxyguanosine (**8-OHDG**), triglycerides (**TG**), D-lactic acid (**D-LA**), total antioxidant capacity (**T-AOC**); and activities of catalase (**CAT**), glutathione peroxidase (**GSH-Px**), and superoxide dismutase (**SOD**). Serum levels of physiological and biochemical parameters and indicators of antioxidant capacity were measured using commercial kits (Jiancheng Bioengineering Institute, Nanjing, China).

### Intestinal Antioxidant Capacity Indicators

To detect antioxidant levels in the duodenum, jejunum, and ileum, T-AOC, CAT, GSH-Px, MDA, and 8-OHDG levels and serum SOD activity were measured using commercial kits (Jiancheng Bioengineering Institute, Nanjing, China).

### Real-time quantitative polymerase chain reaction (PCR)

Total RNA was extracted from duodenal, jejunal, and ileal tissue samples using the Trizol reagent (TransGen Biotech, Beijing, China) according to the manufacturer’s instructions. RNA integrity was assessed by visualization using agarose gels. RNA concentration and purity were determined using a Nanodrop Gentier 48e spectrophotometer (Laboao Scientific, Zhengzhou, China). cDNA synthesis was performed according to the manufacturer’s instructions (TransGen Biotech, Nanjing, China) using primers based on the geese genome sequence (Table 2) synthesized by Sagon Biotech (Shanghai, China). Reverse transcription was performed for 5 min at 50°C and terminated for 5 s at 85°C. The gene for 18S was used as a reference for normalization. Quantitative real-time PCR was performed on a CFX Connect Real-Time PCR Detection System (Thermo Fisher Scientific., MA, USA) using an SYBR Green PCR kit (TransGen Biotech, Beijing, China). The real-time PCR program was initiated with denaturation at 94°C for 30 s, followed by 40 cycles at 94°C for 5 s and 60°C for 30 s. Dissociation analysis of the amplification products was performed after each PCR run to confirm that a single PCR product was amplified and detected. Results were calculated using the 2^−ΔΔCT^ method

**Table 2.**
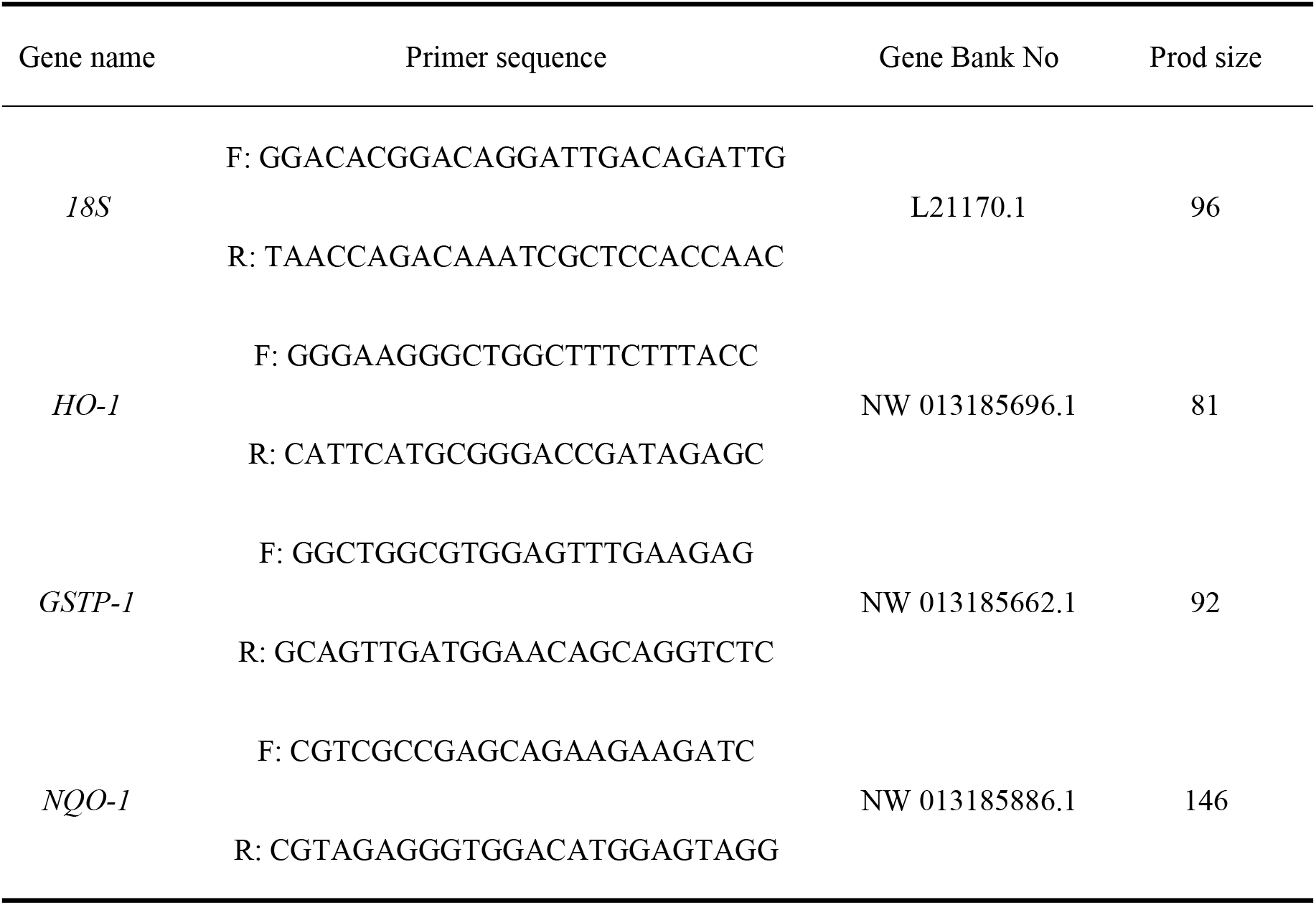
Real-time PCR primer sequences.

#### Data Analysis

The results are presented as the mean and pooled SEM. Data are analyzed by one-way ANOVA using SPSS statistical software (version 26.0, SPSS, IBM Corp., Armonk, NY, USA). Duncan’s multiple range test was used to determine significant differences among treatment means. Differences were considered statistically significant at *P* < 0.05.

## Results

### Growth Performance

Table 3 presents the effects of dietary quercetin supplementation on growth performance indexes. Considering d 1 to 16, group Q2 exhibited a significantly higher FBW value than groups C and L (*P <* 0.05). Considering d 17 to 21, groups Q1 and Q4 had significantly lower FBW values than group C (*P <* 0.05), groups Q2 and Q3 presented significantly higher values than group L, while groups Q1 and Q4 had significantly lower values than group C (*P <* 0.05). The FCR of quercetin-supplemented groups was significantly reduced at d 17 to 21 when compared with that of group L (*P <* 0.05). Considering d 1 to 21, ADG values of groups Q1, Q2, and Q3 were significantly higher than those in group L but significantly lower than those of group C (*P <* 0.05). In addition, the ADG values of group Q4 were significantly lower than that of group C (*P <* 0.05). Groups Q1, Q2, and Q3 showed significantly lower ADG values than group L (*P <* 0.05).

**Table 3.**
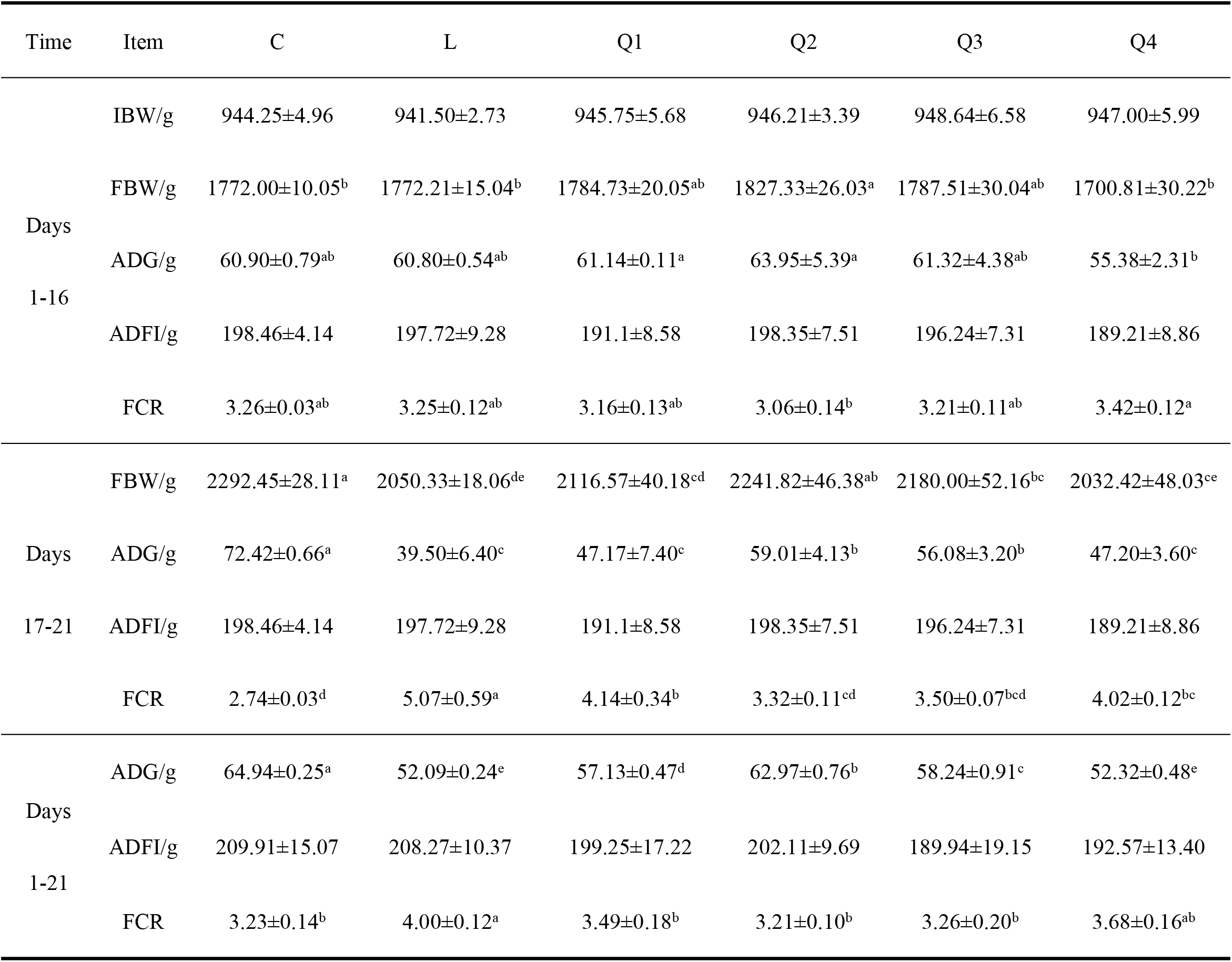
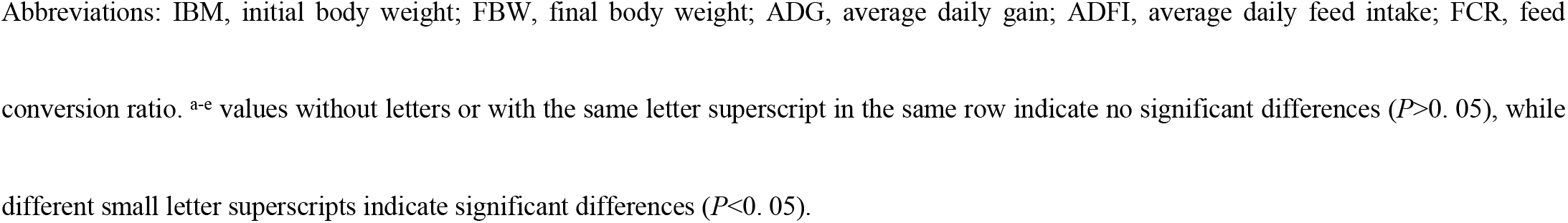
Effects of dietary quercetin supplementation on growth performance indexes in Jilin White geese.

### Serum Physiological and Biochemical Parameters

As shown in Table 4, group L had a significantly lower HDL-C value than group C (*P* < 0.05). LDL-C in groups Q2 and Q3 was significantly lower than that in group L (*P* < 0.05), and group Q1 showed a significantly lower value than group C (*P* < 0.05). Compared with group L, groups who received dietary quercetin supplementation had significantly reduced serum AST concentrations (*P* < 0.05). The ALT level of group Q4 was lower than that of group L (*P* < 0.05). Furthermore, groups Q1, Q2, and Q3 had significantly higher ALT values than group C (*P* < 0.05). In addition, TG levels were significantly lower in group Q4 than those in group C (*P* < 0.05).

**Table 4.**
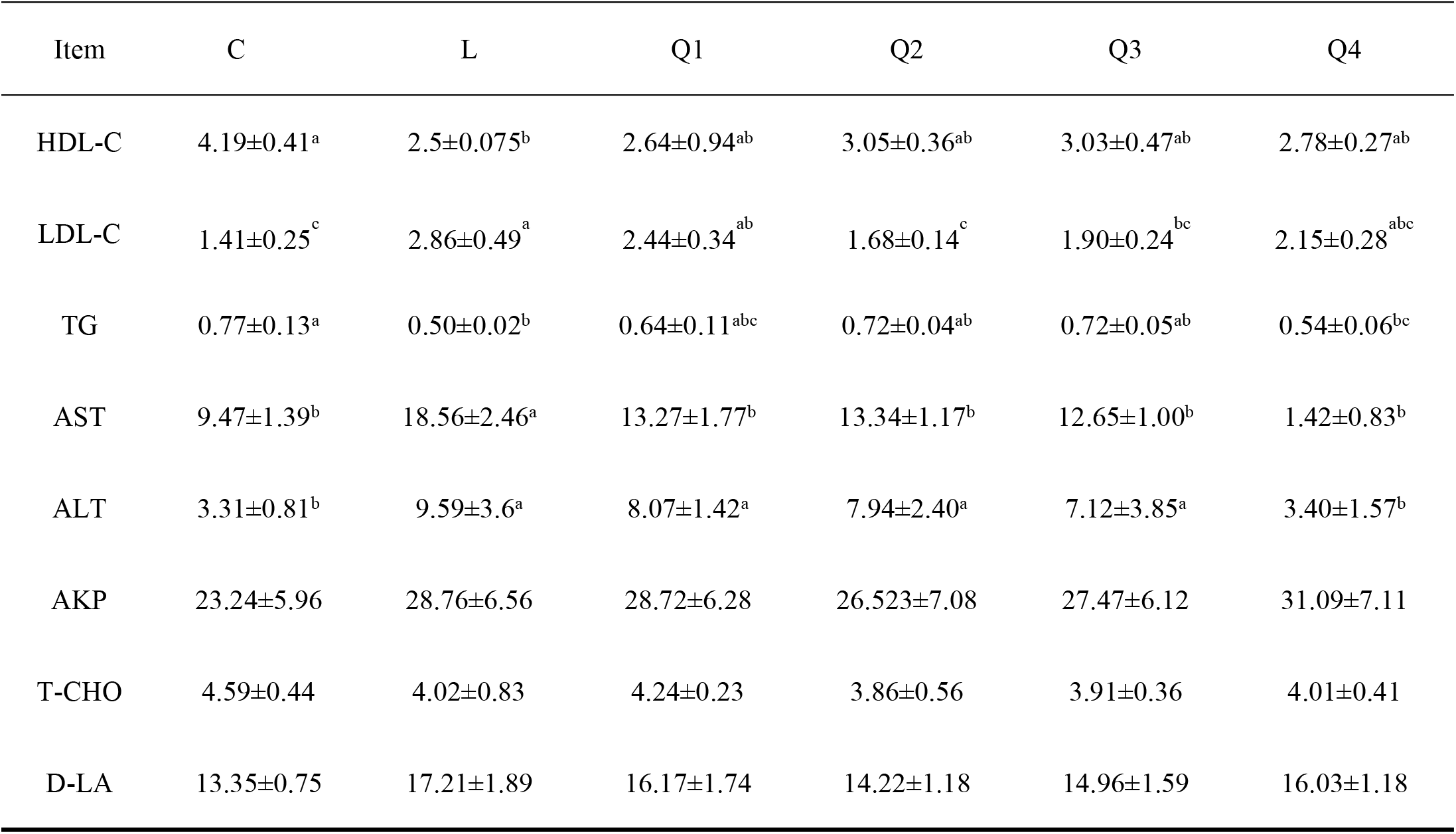
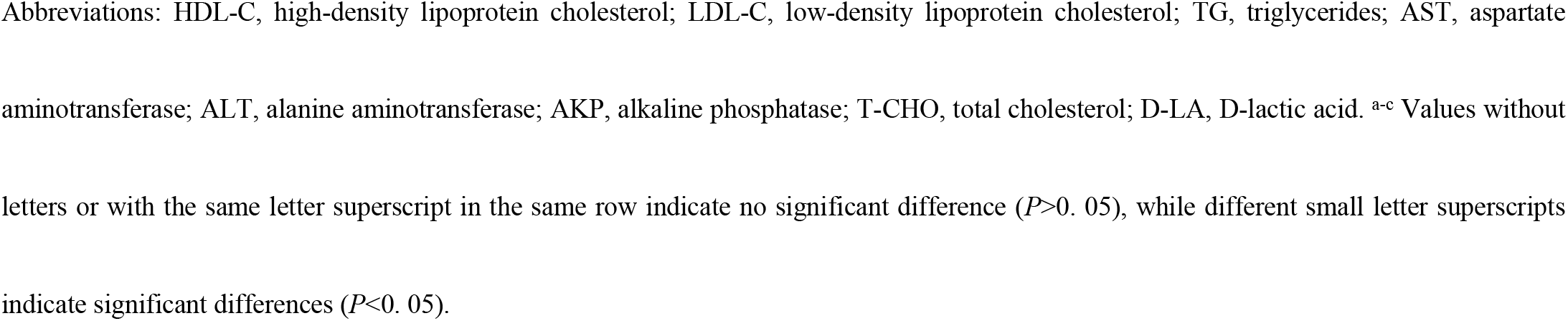
Effects of dietary quercetin supplementation on serum physiology and biochemistry indexes in Jilin White geese.

| Item | C | L | Q1 | Q2 | Q3 | Q4 |
| --- | --- | --- | --- | --- | --- | --- |
| HDL-C | 4.19±0.41<sup>a</sup> | 2.5±0.075<sup>b</sup> | 2.64±0.94<sup>ab</sup> | 3.05±0.36<sup>ab</sup> | 3.03±0.47<sup>ab</sup> | 2.78±0.27<sup>ab</sup> |
| LDL-C | 1.41±0.25<sup>c</sup> | 2.86±0.49<sup>a</sup> | 2.44±0.34<sup>ab</sup> | 1.68±0.14<sup>c</sup> | 1.90±0.24<sup>bc</sup> | 2.15±0.28<sup>abc</sup> |
| TG | 0.77±0.13<sup>a</sup> | 0.50±0.02<sup>b</sup> | 0.64±0.11<sup>abc</sup> | 0.72±0.04<sup>ab</sup> | 0.72±0.05<sup>ab</sup> | 0.54±0.06<sup>bc</sup> |
| AST | 9.47±1.39<sup>b</sup> | 18.56±2.46<sup>a</sup> | 13.27±1.77<sup>b</sup> | 13.34±1.17<sup>b</sup> | 12.65±1.00<sup>b</sup> | 1.42±0.83<sup>b</sup> |
| ALT | 3.31±0.81<sup>b</sup> | 9.59±3.6<sup>a</sup> | 8.07±1.42<sup>a</sup> | 7.94±2.40<sup>a</sup> | 7.12±3.85<sup>a</sup> | 3.40±1.57<sup>b</sup> |
| AKP | 23.24±5.96 | 28.76±6.56 | 28.72±6.28 | 26.523±7.08 | 27.47±6.12 | 31.09±7.11 |
| T-CHO | 4.59±0.44 | 4.02±0.83 | 4.24±0.23 | 3.86±0.56 | 3.91±0.36 | 4.01±0.41 |
| D-LA | 13.35±0.75 | 17.21±1.89 | 16.17±1.74 | 14.22±1.18 | 14.96±1.59 | 16.03±1.18 |

Abbreviations: HDL-C, high-density lipoprotein cholesterol; LDL-C, low-density lipoprotein cholesterol; TG, triglycerides; AST, aspartate aminotransferase; ALT, alanine aminotransferase; AKP, alkaline phosphatase; T-CHO, total cholesterol; D-LA, D-lactic acid. <sup>a-c</sup> Values without letters or with the same letter superscript in the same row indicate no significant difference ( $P>0.05$ ), while different small letter superscripts indicate significant differences ( $P<0.05$ ).

### Serum Antioxidant Capacity

Table 5 illustrates the effects of dietary quercetin supplementation on serum antioxidant capacity. Compared with group L, groups subjected to dietary quercetin supplementation had significantly increased T-AOC and CAT values (*P <* 0.05). The GSH-Px activity of group C was significantly higher than that of group L (*P* < 0.05). Quercetin-supplemented groups had higher levels of 8-OHDG than group C (*P* < 0.05). Compared with group L, quercetin-supplemented groups showed significantly reduced MDA concentrations (*P* < 0.05). The SOD activity in groups Q2 and Q3 was significantly higher than in group L (*P* < 0.05).

**Table 5.**
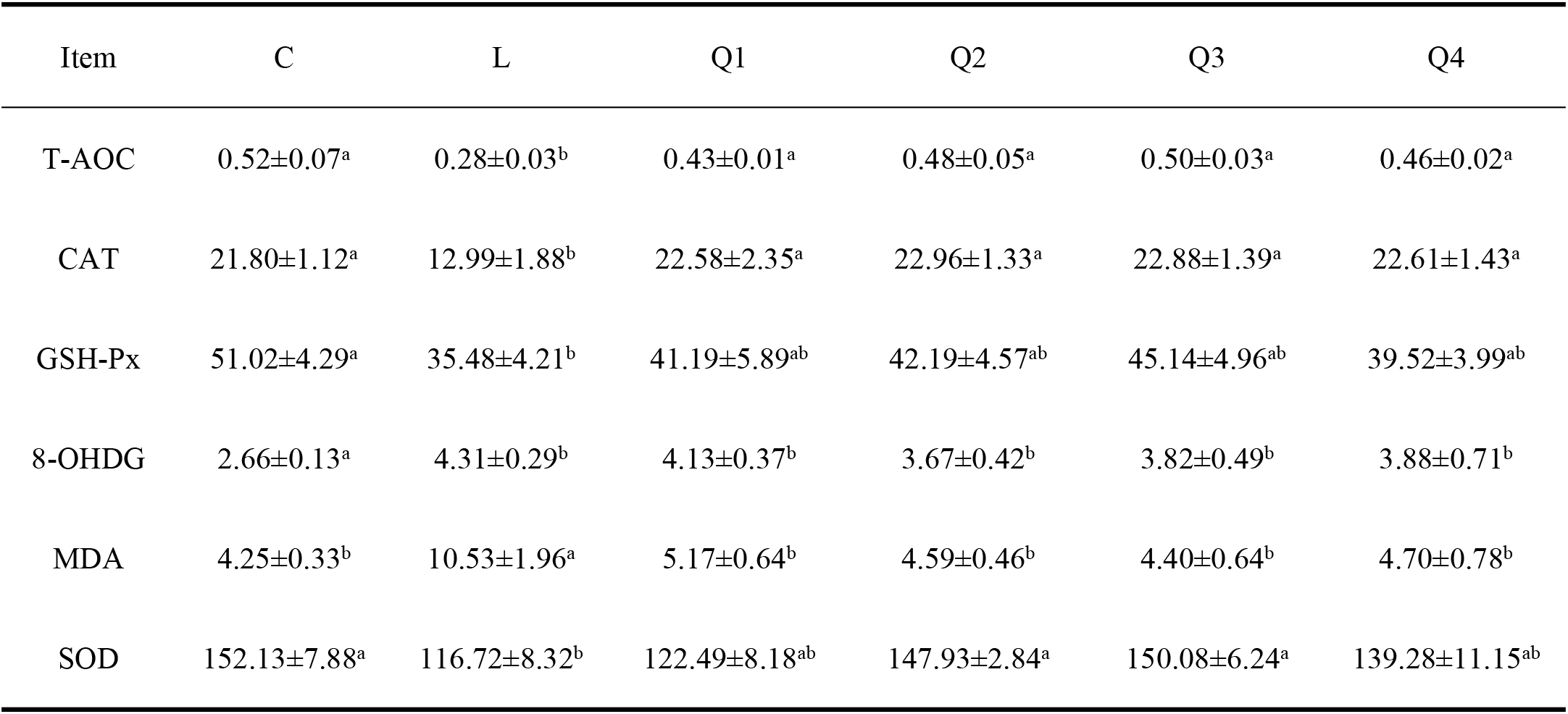
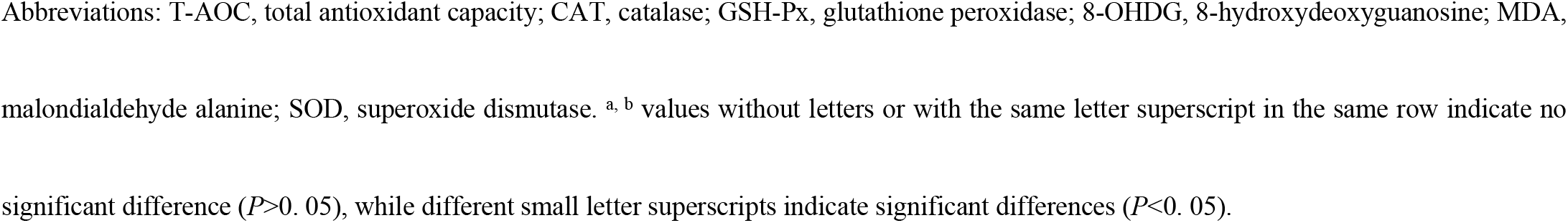
Effects of dietary quercetin supplementation on serum antioxidant capacity of Jilin White geese.

| Item | C | L | Q1 | Q2 | Q3 | Q4 |
| --- | --- | --- | --- | --- | --- | --- |
| T-AOC | 0.52±0.07<sup>a</sup> | 0.28±0.03<sup>b</sup> | 0.43±0.01<sup>a</sup> | 0.48±0.05<sup>a</sup> | 0.50±0.03<sup>a</sup> | 0.46±0.02<sup>a</sup> |
| CAT | 21.80±1.12<sup>a</sup> | 12.99±1.88<sup>b</sup> | 22.58±2.35<sup>a</sup> | 22.96±1.33<sup>a</sup> | 22.88±1.39<sup>a</sup> | 22.61±1.43<sup>a</sup> |
| GSH-Px | 51.02±4.29<sup>a</sup> | 35.48±4.21<sup>b</sup> | 41.19±5.89<sup>ab</sup> | 42.19±4.57<sup>ab</sup> | 45.14±4.96<sup>ab</sup> | 39.52±3.99<sup>ab</sup> |
| 8-OHDG | 2.66±0.13<sup>a</sup> | 4.31±0.29<sup>b</sup> | 4.13±0.37<sup>b</sup> | 3.67±0.42<sup>b</sup> | 3.82±0.49<sup>b</sup> | 3.88±0.71<sup>b</sup> |
| MDA | 4.25±0.33<sup>b</sup> | 10.53±1.96<sup>a</sup> | 5.17±0.64<sup>b</sup> | 4.59±0.46<sup>b</sup> | 4.40±0.64<sup>b</sup> | 4.70±0.78<sup>b</sup> |
| SOD | 152.13±7.88<sup>a</sup> | 116.72±8.32<sup>b</sup> | 122.49±8.18<sup>ab</sup> | 147.93±2.84<sup>a</sup> | 150.08±6.24<sup>a</sup> | 139.28±11.15<sup>ab</sup> |

Abbreviations: T-AOC, total antioxidant capacity; CAT, catalase; GSH-Px, glutathione peroxidase; 8-OHDG, 8-hydroxydeoxyguanosine; MDA, malondialdehyde alanine; SOD, superoxide dismutase. <sup>a, b</sup> values without letters or with the same letter superscript in the same row indicate no significant difference ( $P>0.05$ ), while different small letter superscripts indicate significant differences ( $P<0.05$ ).

### Antioxidant Capacity of the Duodenum

Table 6 illustrates the effects of dietary quercetin supplementation on the duodenal antioxidant capacity. Groups Q1, Q2, and Q3 had significantly higher T-AOC levels than group L (*P* < 0.05). Groups Q1 and Q4 had significantly lower T-AOC levels than group C (*P <* 0.05). The CAT activity of groups Q1, Q2, and Q4 was significantly lower than that of group C (*P <* 0.05), whereas group Q3 exhibited significantly higher activity than group L (*P <* 0.05). Group C had significantly higher GSH-Px and SOD activities than group L (*P <* 0.05). 8-OHDG levels of groups Q2 and Q3 were significantly lower than those of group L (*P <* 0.05). Groups Q2, Q3, and Q4 exhibited significantly lower MDA values than group L (*P <* 0.05), whereas group Q1 had a significantly higher value than group C (*P <* 0.05).

**Table 6.**
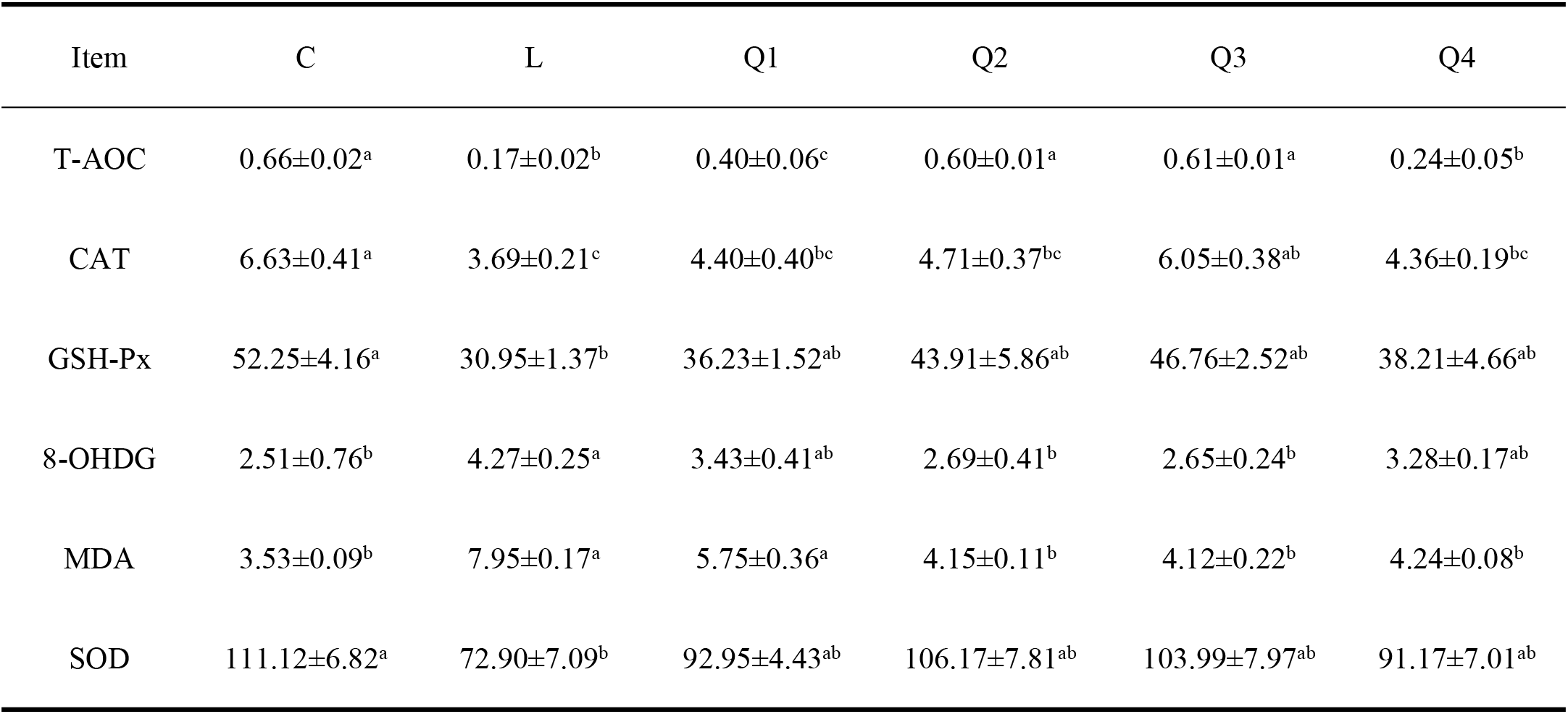
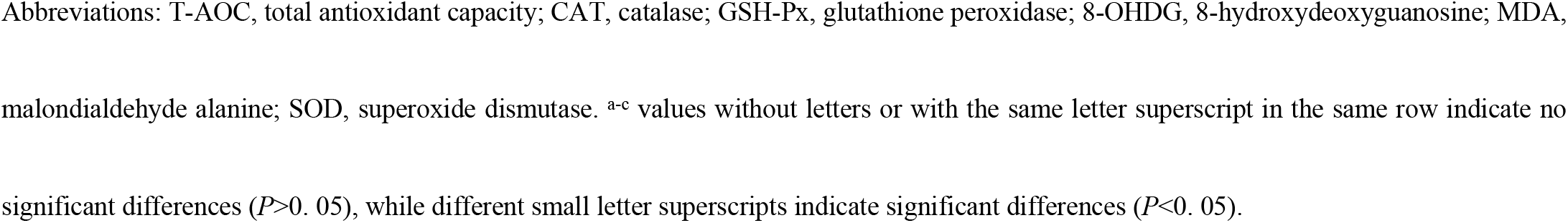
Effects of dietary quercetin supplementation on the duodenal antioxidant capacity of Jilin White geese.

| Item | C | L | Q1 | Q2 | Q3 | Q4 |
| --- | --- | --- | --- | --- | --- | --- |
| T-AOC | 0.66±0.02<sup>a</sup> | 0.17±0.02<sup>b</sup> | 0.40±0.06<sup>c</sup> | 0.60±0.01<sup>a</sup> | 0.61±0.01<sup>a</sup> | 0.24±0.05<sup>b</sup> |
| CAT | 6.63±0.41<sup>a</sup> | 3.69±0.21<sup>c</sup> | 4.40±0.40<sup>bc</sup> | 4.71±0.37<sup>bc</sup> | 6.05±0.38<sup>ab</sup> | 4.36±0.19<sup>bc</sup> |
| GSH-Px | 52.25±4.16<sup>a</sup> | 30.95±1.37<sup>b</sup> | 36.23±1.52<sup>ab</sup> | 43.91±5.86<sup>ab</sup> | 46.76±2.52<sup>ab</sup> | 38.21±4.66<sup>ab</sup> |
| 8-OHDG | 2.51±0.76<sup>b</sup> | 4.27±0.25<sup>a</sup> | 3.43±0.41<sup>ab</sup> | 2.69±0.41<sup>b</sup> | 2.65±0.24<sup>b</sup> | 3.28±0.17<sup>ab</sup> |
| MDA | 3.53±0.09<sup>b</sup> | 7.95±0.17<sup>a</sup> | 5.75±0.36<sup>a</sup> | 4.15±0.11<sup>b</sup> | 4.12±0.22<sup>b</sup> | 4.24±0.08<sup>b</sup> |
| SOD | 111.12±6.82<sup>a</sup> | 72.90±7.09<sup>b</sup> | 92.95±4.43<sup>ab</sup> | 106.17±7.81<sup>ab</sup> | 103.99±7.97<sup>ab</sup> | 91.17±7.01<sup>ab</sup> |

Abbreviations: T-AOC, total antioxidant capacity; CAT, catalase; GSH-Px, glutathione peroxidase; 8-OHDG, 8-hydroxydeoxyguanosine; MDA, malondialdehyde alanine; SOD, superoxide dismutase. <sup>a-c</sup> values without letters or with the same letter superscript in the same row indicate no significant differences ( $P>0.05$ ), while different small letter superscripts indicate significant differences ( $P<0.05$ ).

### Antioxidant Capacity of the Jejunum

Table 7 summarizes the effects of dietary quercetin supplementation on the jejunal antioxidant capacity. The AOC level and CAT activity of group Q2 h were significantly higher than those of group L (P < 0.05). The GSH-Px and SOD values of group C were significantly higher than those of group L (P < 0.05). Groups Q2 and Q3 had significantly lower 8-OHDG levels than group L (P < 0.05). Group Q4 had a significantly higher 8-OHDG level than group C (P < 0.05).

**Table 7.** Effects of dietary quercetin supplementation on the jejunal antioxidant capacity of Jilin White geese.

| Item | C | L | Q1 | Q2 | Q3 | Q4 |
| --- | --- | --- | --- | --- | --- | --- |
| AOC | 0.62±0.08<sup>a</sup> | 0.29±0.09<sup>b</sup> | 0.47±0.06<sup>ab</sup> | 0.53±0.04<sup>ab</sup> | 0.62±0.09<sup>a</sup> | 0.44±0.04<sup>ab</sup> |
| CAT | 6.63±0.41<sup>a</sup> | 3.69±0.21<sup>c</sup> | 4.40±0.40<sup>bc</sup> | 4.71±0.37<sup>bc</sup> | 6.05±0.38<sup>ab</sup> | 4.36±0.19<sup>bc</sup> |
| GSH-Px | 41.56±6.63<sup>a</sup> | 27.57±5.35<sup>b</sup> | 30.33±3.82<sup>ab</sup> | 36.67±3.31<sup>ab</sup> | 39.17±4.11<sup>ab</sup> | 33.47±7.96<sup>ab</sup> |
| 8-OHDG | 3.23±0.16<sup>a</sup> | 5.59±0.73<sup>c</sup> | 5.48±0.67<sup>c</sup> | 3.75±0.47<sup>ab</sup> | 3.53±0.41<sup>ab</sup> | 4.86±0.12<sup>bc</sup> |
| MDA | 21.81±5.16<sup>b</sup> | 37.13±2.97<sup>a</sup> | 29.2±9.80<sup>ab</sup> | 26.6±1.97<sup>ab</sup> | 27.81±3.89<sup>ab</sup> | 35.6±1.80<sup>a</sup> |
| SOD | 110.23±3.48<sup>b</sup> | 56.63±4.60<sup>a</sup> | 74.83±7.60<sup>ab</sup> | 93.33±5.9<sup>ab</sup> | 96.29±7.96<sup>ab</sup> | 65.99±3.95<sup>ab</sup> |

Abbreviations: T-AOC, total antioxidant capacity; CAT, catalase; GSH-Px, glutathione peroxidase; 8-OHDG, 8-hydroxydeoxyguanosine; MDA, malondialdehyde alanine; SOD, superoxide dismutase. <sup>a-c</sup> Values without letters or with the same letter superscript in the same row indicate no significant difference ( $P>0.05$ ), while different small letter superscripts indicate significant differences ( $P<0.05$ ).

### Antioxidant Capacity of the Ileum

As shown in Table 8, quercetin-supplemented groups had significantly higher AOC levels than group L (*P* < 0.05). CAT activities of groups Q2 and Q3 were significantly higher than those of group L (*P* < 0.05). Group Q1 had significantly lower GSH-Px activity than group L (*P* < 0.05). Groups Q2 and Q3 had significantly lower MDA values than group L, and group Q3 had significantly higher SOD activity than group L (*P* < 0.05).

**Table 8.**
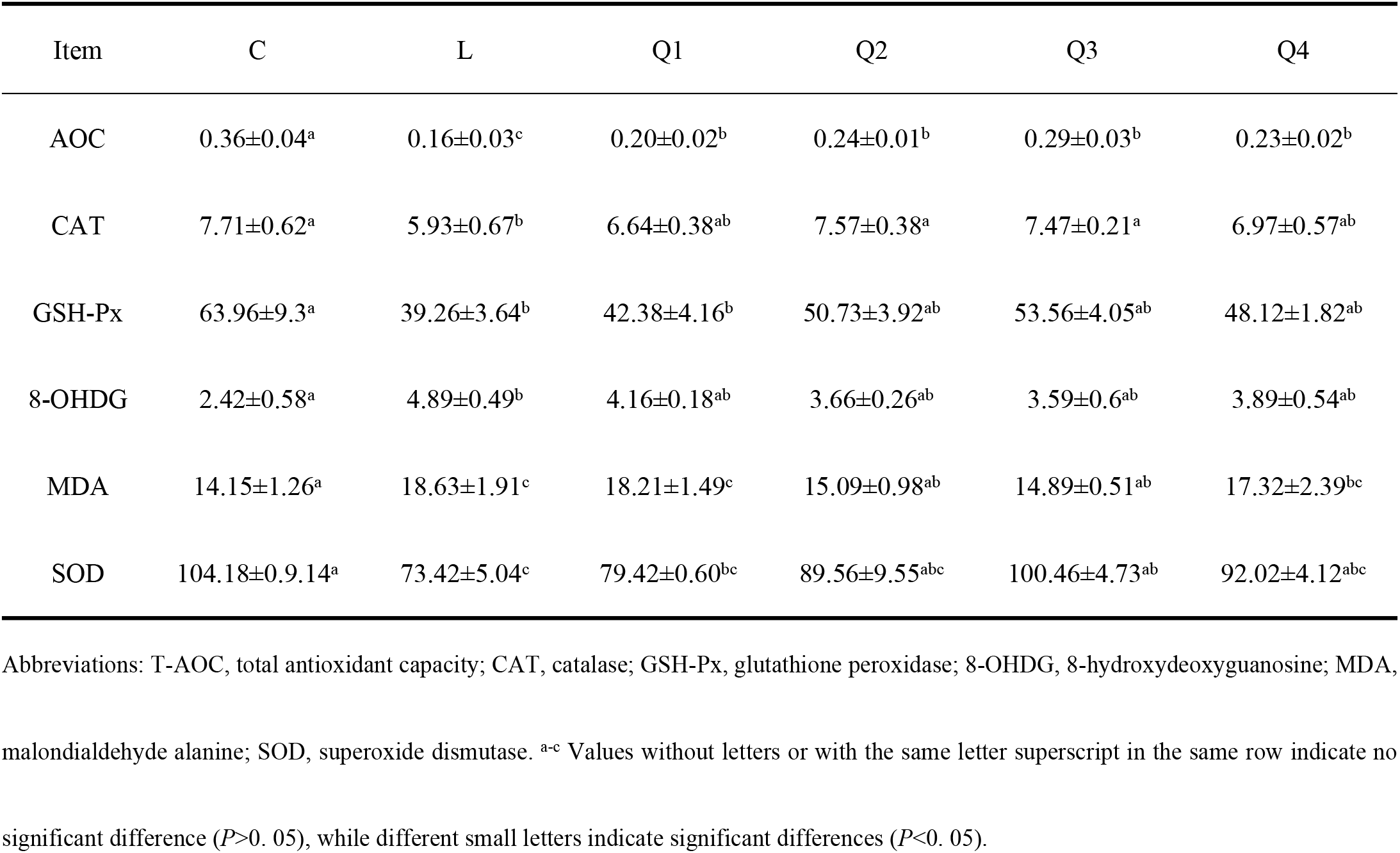
Effects of quercetin on the ileal antioxidant capacity of Jilin White geese.

| Item | C | L | Q1 | Q2 | Q3 | Q4 |
| --- | --- | --- | --- | --- | --- | --- |
| AOC | 0.36±0.04<sup>a</sup> | 0.16±0.03<sup>c</sup> | 0.20±0.02<sup>b</sup> | 0.24±0.01<sup>b</sup> | 0.29±0.03<sup>b</sup> | 0.23±0.02<sup>b</sup> |
| CAT | 7.71±0.62<sup>a</sup> | 5.93±0.67<sup>b</sup> | 6.64±0.38<sup>ab</sup> | 7.57±0.38<sup>a</sup> | 7.47±0.21<sup>a</sup> | 6.97±0.57<sup>ab</sup> |
| GSH-Px | 63.96±9.3<sup>a</sup> | 39.26±3.64<sup>b</sup> | 42.38±4.16<sup>b</sup> | 50.73±3.92<sup>ab</sup> | 53.56±4.05<sup>ab</sup> | 48.12±1.82<sup>ab</sup> |
| 8-OHDG | 2.42±0.58<sup>a</sup> | 4.89±0.49<sup>b</sup> | 4.16±0.18<sup>ab</sup> | 3.66±0.26<sup>ab</sup> | 3.59±0.6<sup>ab</sup> | 3.89±0.54<sup>ab</sup> |
| MDA | 14.15±1.26<sup>a</sup> | 18.63±1.91<sup>c</sup> | 18.21±1.49<sup>c</sup> | 15.09±0.98<sup>ab</sup> | 14.89±0.51<sup>ab</sup> | 17.32±2.39<sup>bc</sup> |
| SOD | 104.18±0.9.14<sup>a</sup> | 73.42±5.04<sup>c</sup> | 79.42±0.60<sup>bc</sup> | 89.56±9.55<sup>abc</sup> | 100.46±4.73<sup>ab</sup> | 92.02±4.12<sup>abc</sup> |

Abbreviations: T-AOC, total antioxidant capacity; CAT, catalase; GSH-Px, glutathione peroxidase; 8-OHDG, 8-hydroxydeoxyguanosine; MDA, malondialdehyde alanine; SOD, superoxide dismutase. <sup>a-c</sup> Values without letters or with the same letter superscript in the same row indicate no significant difference ( $P > 0.05$ ), while different small letters indicate significant differences ( $P < 0.05$ ).

### Effects of Quercetin Supplementation on the Expression of Genes Involved in Oxidative Stress in the Duodenum

As shown in Figure 1, duodenal expression of *HO-1* and *NQO-1* mRNA was significantly decreased in quercetin-supplemented groups treated, intraperitoneally administered LPS, when compared with that in group C (*P <* 0.01; Figure 1A and C). *HO-1* mRNA expression was significantly higher in group Q3 than that in group L (*P <* 0.01; Figure 1A). *NQO-1* mRNA expression was significantly higher in groups Q3 and Q4 than that in group L (*P <* 0.01; Figure 1C). Compared with group C, groups L and Q1 had significantly reduced *GSTP* mRNA expression (*P <* 0.01); this expression level was significantly decreased in groups Q2 and Q4. Groups Q2 and Q4 had significantly reduced GSTP mRNA expression when compared with that in group L(*P <* 0.05). Expression of *GSTP* mRNA was significantly lower in group Q3 than that in group L (*P <* 0.01; Figure 1B).

**Figure 1.**
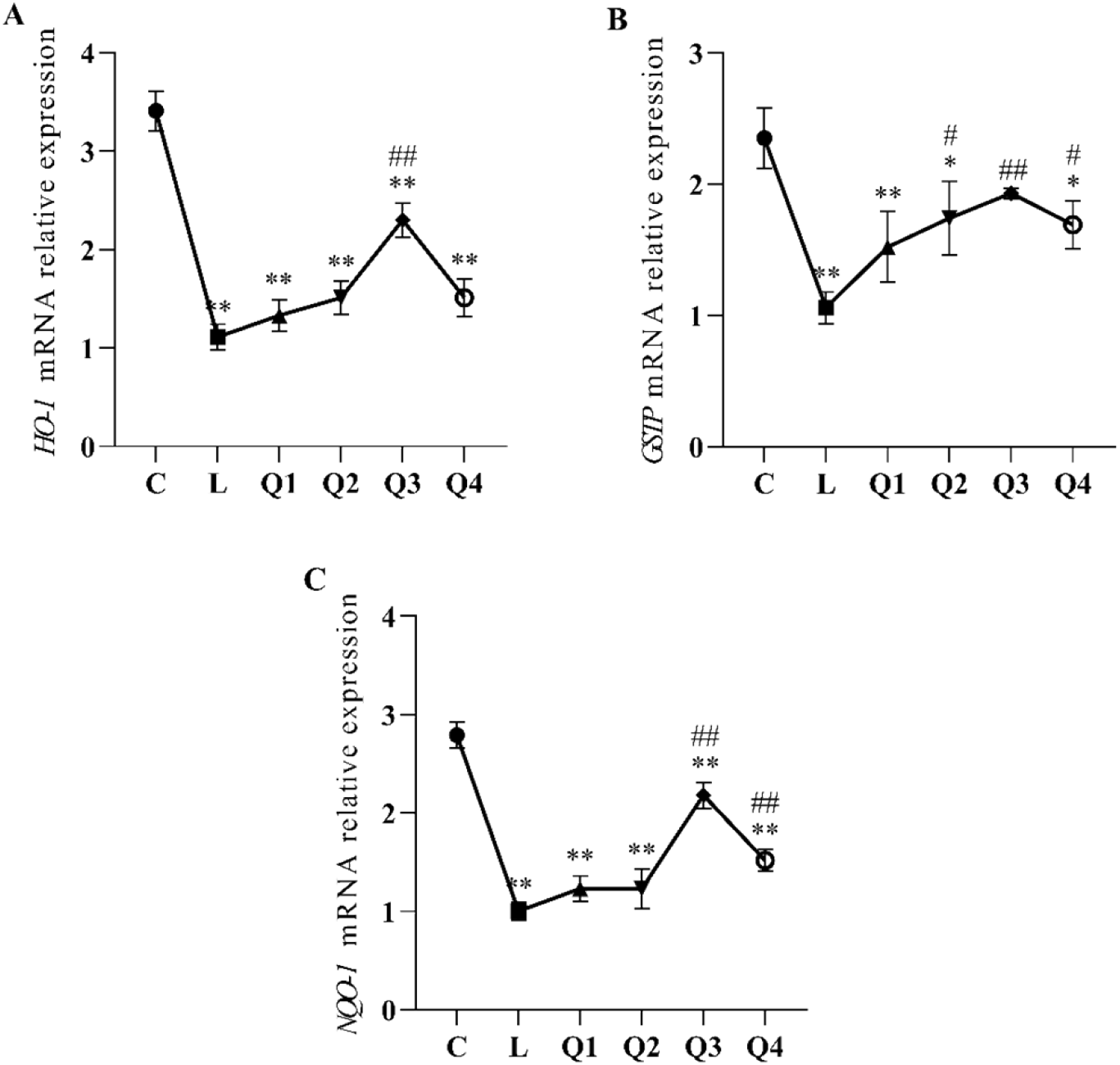
Effects of dietary quercetin supplementation on mRNA levels of HO-1, GSTP, and NQO-1 in the geese duodenum after LPS injection (n = 8). Relative gene expression of HO-1 (A), GSTP (B), and NQO-1 (C) in the duodenum after LPS injection. The gene for 18S was used as a reference for normalization. * Indicates significant difference compared with (P<0.05). ** Indicates extremely significant difference compared with group C (P<0.01). # Indicates significant difference compared with group L (P<0.05). ## Indicates extremely significant difference compared with group L (P<0.01). HO-1, heme oxygenase 1; NQO1, reduced coenzyme/quinone oxidoreductase; GSTP, glutathione oxidase; LPS, lipopolysaccharide.

### Effects of Quercetin Supplementation on the Expression of Genes Involved in Oxidative Stress in the Jejunum

Figure 2 illustrates the effects of quercetin supplementation on gene expression in the jejunum following the induction of oxidative stress. Compared with group C, expression of *HO-1* mRNA was significantly reduced in quercetin-supplemented groups administered i.p. LPS (*P <* 0.01). *HO-1* mRNA expression was significantly lower in group Q3 than that in group L (*P <* 0.01; Figure 2A). Groups Q1, Q2, and Q4 exhibited significantly reduced *GSTP* mRNA expression after i.p. LPS administration when compared with that in group C (*P <* 0.01). Group Q3 had significantly lower *GSTP* mRNA expression than groups C and L (*P <* 0.05; Figure 2B). Compared with group C, the expression of *NQO-1* mRNA was significantly reduced in groups L, Q1 and Q4 (*P <* 0.01). Groups Q1 and Q4 had higher *NQO-1* mRNA expression than group L, but the difference was not statistically significant (*P >* 0.05). Furthermore, groups Q2 and Q3 showed significantly higher *NQO-1* mRNA expression than group L (*P <* 0.01; Figure 2C).

**Figure 2.**
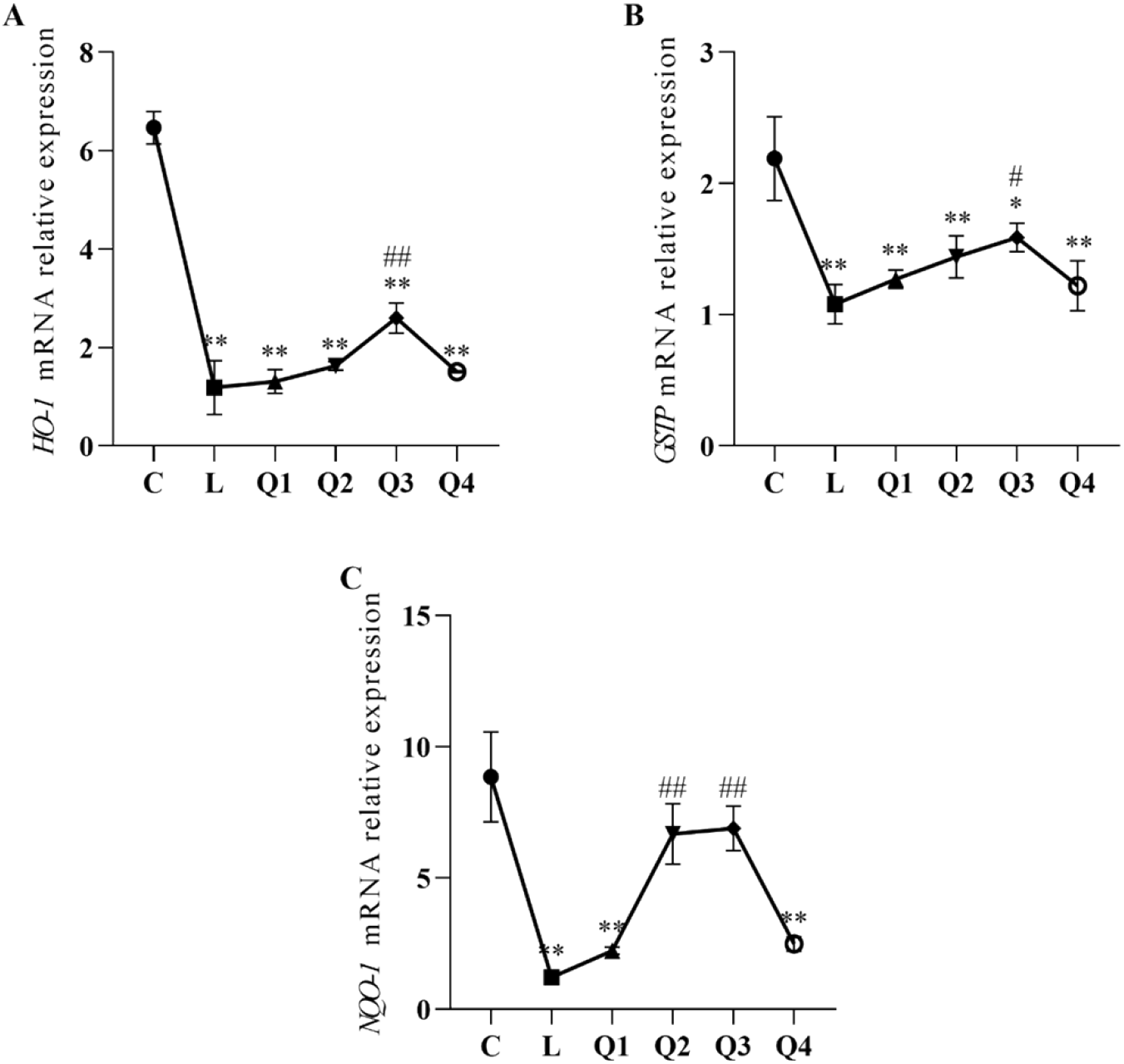
Effects of dietary quercetin supplementation on mRNA levels of HO-1, GSTP, and NQO-1 in the geese jejunum after LPS injection (n = 8). Relative gene expression of HO-1 (A), GSTP (B), and NQO-1 (C) in the jejunum after LPS injection. The gene for 18S was used as a reference for normalization. * Indicates significant difference compared with group C (P<0.05). ** Indicates extremely significant difference compared with group C (P<0.01). # Indicates significant difference compared with group L (P<0.05). ## Indicates extremely significant difference compared with group L (P<0.01). HO-1, heme oxygenase 1; NQO1, reduced coenzyme/quinone oxidoreductase; GSTP, glutathione oxidase; LPS, lipopolysaccharide.

### Effects of Quercetin Supplementation on the Expression of Genes Involved in Oxidative Stress in the Ileum

Figure 3 illustrates the effects of quercetin supplementation on gene expression in the ileum following the induction of oxidative stress. Compared with group C, *HO-1* mRNA expression was significantly reduced in groups L, Q1, and Q4 (*P <* 0.01). *HO-1* mRNA expression was significantly higher in group Q3 than that in group L (*P <* 0.01), while that in group Q2 was significantly higher than that in group L (*P <* 0.05) (Figure 3A). The expression of *GSTP* mRNA was significantly reduced in groups L, Q1, and Q4 when compared with that in group C (*P <* 0.01). Groups Q2 and Q3 showed significantly lower *GSTP* mRNA expression than group C (*P <* 0.05), while group Q3 had significantly lower expression than group L (*P >* 0.05) (Figure 3B). Expression of *NQO-1* mRNA was significantly reduced in quercetin-supplemented groups administered i.p. LPS when compared with that in group C (*P <* 0.01). Furthermore, groups Q2 and Q3 had significantly lower *NQO-1* mRNA expression than group L (*P <* 0.01), while group Q1 exhibited significantly lower expression than group L (*P <* 0.01) (Figure 3C).

**Figure 3.**
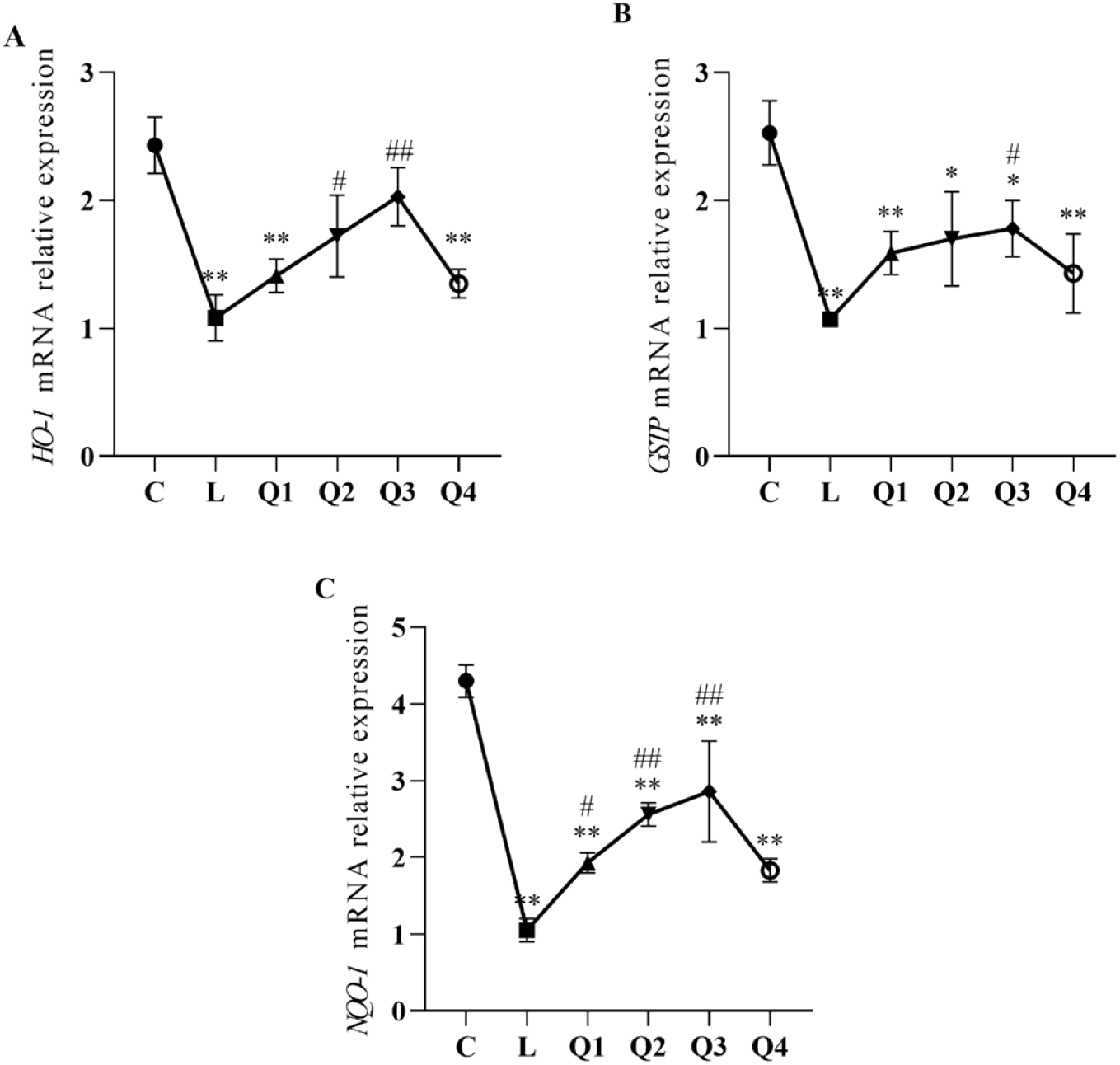
Effects of dietary quercetin supplementation on mRNA levels of HO-1, GSTP, and NQO-1 in the geese ileum after LPS injection (n = 8). Relative expression of the gene for HO-1 (A), GSTP (B), and NQO-1 (C) in the ileum after LPS injection. The gene for 18S was used as a reference for normalization. * Indicates significant difference compared with group C (P<0.05). ** Indicates extremely significant differences compared with group C (P<0.01). # Indicates significant difference compared with group L (P<0.05). ## Indicates extremely significant difference compared with group L (P<0.01). HO-1, heme oxygenase 1; NQO1, reduced coenzyme/quinone oxidoreductase; GSTP, glutathione oxidase; LPS, lipopolysaccharide.

## Discussion

Oxidative stress can impair growth performance and productivity and negatively impact animal health [13,14]. Dietary supplementation with quercetin was shown to increase egg production in hens [15]. Adding quercetin to poultry diets reduced the effects of oxidative stress on animal performance [16]. In the present study, incorporating 300 mg/kg quercetin into the diet could increase the FBW of Jilin White geese before oxidative stress. Adding 300 or 450 mg/kg of quercetin to the diet improved the performance of Jilin White geese after oxidative stress.

Herein, blood levels of MDA and 8-OHDG in Jilin White geese were significantly higher in group L than those in the control group, while the AOC level and CAT, SOD, and GSH-Px activities were significantly lower. These results indicate that the geese experienced oxidative damage and that an oxidative stress model was successfully established.

T-CHO and TG are important indicators of lipid metabolism in the body. T-CHO is synthesized and stored in the liver, and the development of adipose tissue in birds and the extent of deposition depends on TG levels [17]. Serum lactate dehydrogenase is used as a plasma marker of intestinal permeability. Quercetin reportedly exerts a considerable hypolipidemic effect [18] and notably reduces serum lactate dehydrogenase [19]. The addition of 300 and 450 mg/kg quercetin to the diet could significantly reduce HDL-C, thereby indicating that quercetin can regulate fat deposition. AST and ALT are mainly found in the liver and are important enzymes that respond to liver function. In the present study, group L had a significantly higher AST level than group C, indicating the presence of oxidative stress-induced liver damage. Adding quercetin to the diet could reduce AST and ALT levels, potentially inhibiting lipid peroxidation and playing a protective role against hepatocyte damage.

Nrf2 is an important regulator of the antioxidant defense system, regulating the expression of antioxidant enzymes and protecting cells from damage caused by oxidative stress [20]. In response to oxidative stress, Nrf2 translocates to the nucleus and interacts with antioxidant response elements to initiate transcription of *SOD, CAT*, and *GSH-Px* [21,22]. Glutathione is synthesized in the liver and is crucial for protecting cells from oxidative stress [23]. Quercetin has been shown to stabilize antioxidant enzymes in rats to normal levels [24]. Herein, dietary supplementation with 300 and 450 mg/kg quercetin could significantly enhance the antioxidant activity in the duodenum, jejunum, and ileum. Quercetin has been shown to reduce dimethylhydrazine-induced colon carcinogenesis in rats via the Nrf2/Keap1-mediated regulation of oxidative DNA damage, DNA repair, and antioxidant system [25]. In the present study, the addition of 450 mg/kg quercetin to the diet of geese significantly upregulated the gene expression of transcription factors *HO-1, GSTP1*, and *NQO1* in the Nrf-2 pathway. These findings suggest that quercetin can upregulate genes related to Nrf2 signaling and thus reduce the damage caused by oxidative stress, which may be achieved by activating genes downstream of the Nrf2 pathway.

## Conclusion

In conclusion, our results revealed that the addition of 300 mg/kg quercetin to the diet under non-stressed conditions could increase the performance of Jilin White geese. Adding 300 mg/kg quercetin to the diet under stressed conditions improved performance after oxidative stress induction, and the AOC level and CAT activity were significantly improved in the duodenum, jejunum, and ileum, effectively reducing oxidative stress-induced damage. The addition of 450 mg/kg quercetin to the diet could inhibit lipid peroxidation and reduce liver cell damage caused by oxidative stress while significantly upregulating the gene expression of transcription factors *HO-1, GSTP1*, and *NQO1* related to the Nrf-2 pathway, thereby alleviating the effect of oxidative stress on intestinal damage in Jilin White geese.

## Author Contributions

**Conceptualization:** Liu Yingkun, Guo Wei, Jia Fangyuan.

**Data curation:** Li Zhichao, Zhang Tao.

**Formal analysis:** Liu Yingkun, Jia Fangyuan.

**Investigation:** Liu Yingkun.

**Methodology:** Wang Kai, Guo Wei, Jia Fangyuan.

**Project administration:** Zhou Haizhu, Guo Wei.

**Result interpretation**: Guo Wei.

**Software:** Liu Mo.

**Supervision:** Zhou Haizhu, Guo Wei.

**Writing – original draft:** Liu Yingkun, Liu Mo

**Writing-review and editing:** Liu Yingkun

